# Flexible binding of m^6^A reader protein YTHDC1 to its preferred RNA motif

**DOI:** 10.1101/787937

**Authors:** Yaozong Li, Rajiv Kumar Bedi, Lars Wiedmer, Danzhi Huang, Paweł Śledź, Amedeo Caflisch

**Affiliations:** Department of Biochemistry, University of Zurich, Winterthurerstrasse 190, CH-8057 Zurich, Switzerland

**Author notes:** To whom correspondence should be addressed., Correspondence may also be addressed to Paweł Śledź.

## Abstract

N6-methyladenosine (m^6^A) is the most prevalent chemical modification in human mRNAs. Its recognition by reader proteins enables many cellular functions, including splicing and translation of mRNAs. However, the binding mechanisms of m^6^A-containing RNAs to their readers are still elusive due to the unclear roles of m^6^A-flanking ribonucleotides. Here, we use a model system, YTHDC1 with its RNA motif 5’-G_−2_G_−1_(m^6^A)C_+1_U_+2_-3’, to investigate the binding mechanisms by atomistic simulations, X-ray crystallography, and isothermal titration calorimetry. The experimental data and simulation results show that m^6^A is captured by an aromatic cage of YTHDC1 and the 3’ terminus nucleotides are stabilized by cation-π-π interactions, while the 5’ terminus remains flexible. Moreover, simulations of unbound RNA motifs reveal that the methyl group of m^6^A and the 5’ terminus shift the conformational preferences of the oligoribonucleotides to the bound-like conformation, thereby facilitating the association process. The proposed binding mechanisms may help in the discovery of chemical probes against m^6^A reader proteins.

## Introduction

Methylation of position N6 in adenines, also known as m^6^A modification, is the most prevalent internal modification of messenger RNA (mRNA) with approximately one in 200 adenine bases modified, and as such it makes significant contributions to central events in biology (Dominissini, Moshitch-Moshkovitz, Salmon-Divon, Amariglio, & Rechavi, 2013; Meyer et al., 2012). On the molecular level, the introduction of m^6^A affects structures of RNAs and their ability to form protein-RNA interactions, and thus modulates processing (Alarcon, Lee, Goodarzi, Halberg, & Tavazoie, 2015; Xiao et al., 2016), translation (Lin, Choe, Du, Triboulet, & Gregory, 2016; X. Wang et al., 2015), and stability of the cellular transcripts (X. Wang et al., 2014). As a consequence, m^6^A is implicated in controlling embryonic development processes and stem cell differentiation (Batista et al., 2014; Geula et al., 2015; Qi et al., 2016; Y. Wang et al., 2014), regulating the mammalian circadian clock (Fustin et al., 2013), and modulating stress response, e.g., heat shock (Zhou et al., 2015). The malfunctions of the cellular machinery regulating the m^6^A modification have in turn been linked to pathologies like obesity (Ben-Haim, Moshitch-Moshkovitz, & Rechavi, 2015), cancer (Peng, Yuan, Jiang, Tang, & Li, 2016), and neurodegeneration (Satterlee et al., 2014). Therefore, the m^6^A field has recently attracted broad attention of drug discovery community (Boriack-Sjodin, Ribich, & Copeland, 2018; Dai, Wang, Zhu, Jin, & Wang, 2018; Delaunay & Frye, 2019; Hsu, Shi, & He, 2017).

While the m^6^A modification has been known for several decades, only recently it has been proven reversible due to the discovery of the selective demethylases (erasers) that remove it from mRNA (Jia et al., 2011; Zheng et al., 2013) and balance the actions of the writer complex (J. Z. Liu et al., 2014; Sledz & Jinek, 2016; P. Wang, Doxtader, & Nam, 2016; X. Wang et al., 2017; Wiedmer, Eberle, Bedi, Sledz, & Caflisch, 2019). Similarly to the epigenetic modifications of histone proteins, the reversibility of the mRNA methylation gives it the potential to constitute an additional layer of epitranscriptomic regulation of gene expression (N. Liu & Pan, 2016; Meyer & Jaffrey, 2014; X. Zhang & Jia, 2016). The reader proteins, i.e., proteins that recognize the modification and elicit response upon it, play the key role at the executive end of such regulation mechanism. A number of proteins have been identified as direct m^6^A readers, with different structures, subcellular localization and mechanism of the recognition of epitranscriptomic mark. There are five YTH-domain-containing m^6^A readers that adopt highly similar open α/β fold and specifically recognize the m^6^A mark through interactions of aromatic cage (F. D. Li, Zhao, Wu, & Shi, 2014; C. Xu et al., 2015; C. Xu et al., 2014; Zhu et al., 2014). Cytoplasmic readers YTHDF1 and YTHDF3 play roles in translation initiation (X. Wang et al., 2015), while highly similar YTHDF2 reader controls destabilization of cellular transcripts by targeting them to P-bodies (X. Wang et al., 2014). The nuclear reader YTHDC1 plays a role in splicing regulation (Xiao et al., 2016), while YTHDC2 contains additional RNAse helicase domain and is important in meiosis (Wojtas et al., 2017).

Specific recognition of m^6^A-containing transcripts by the YTH-domain readers has been studies in details by the means of protein crystallography, pointing out at rigid and conserved recognition of consensus sequence GG(m^6^A)CU. Beyond YTH-domain m^6^A readers, there are also indirect readers (i.e., proteins not interacting with m^6^A but binding to m^6^A-containing transcripts) including HBRNPC (N. Liu et al., 2015), and anti-readers like G3BP1 that bind preferably to unmodified transcripts (Vermeulen, 2017). The crucial role of m^6^A in the molecular recognition has been confirmed, where an aromatic cage consisting of number of key tryptophan residues enables binding. However, the exact role of m^6^A-flanking nucleotides that make a number of hydrogen bonds and salt bridges to the protein (S. K. Luo & Tong, 2014; C. Xu et al., 2014), remains elusive. Further studies on these interactions are crucial not only for understanding the mechanism and timescale of epitranscriptomic regulation, but also for informing the development of therapeutic agents.

It is not easy to infer the importance of the interactions between the m^6^A-flanking nucleotides and reader proteins, partially because they are observed on protein surface and thus may be stabilized by crystal packing. In addition, unbound oligoribonucleotides are highly flexible and may adopt many meta-stable conformations in aqueous solution, which is difficult to study by experiments. To this end, molecular dynamics (MD) may provide complementary information for understanding the binding mechanisms (Dror, Dirks, Grossman, Xu, & Shaw, 2012; Karplus & McCammon, 2002; Sledz & Caflisch, 2018). However, the field of MD simulations on RNA and its complexes with proteins, e.g., m^6^A reader proteins and its ligand RNAs, is less mature than those on proteins (or with organic molecules). In particular, three potential issues hinder the use of RNA-related force fields (Krepl et al., 2015; MacKerell & Nilsson, 2008; Sponer et al., 2017; Vangaveti, Ranganathan, & Chen, 2017): I: the inaccuracy of ribonucleotide force fields themselves (Krepl et al., 2015; Sponer et al., 2017), II: lack of parameters for modified nucleotides (Aduri et al., 2007; Y. Xu, Vanommeslaeghe, Aleksandrov, MacKerell, & Nilsson, 2016) and III: the difficulties in balancing interactions between proteins and RNA molecules (Krepl et al., 2015; MacKerell & Nilsson, 2008).

The first issue has been moderately alleviated by the recent improvements of common RNA force fields, e.g., CHARMM (Denning, Priyakumar, Nilsson, & Mackerell, 2011), AMBER (Tan, Piana, Dirks, & Shaw, 2018) and OPLS (Robertson, Qian, Robinson, Tirado-Rives, & Jorgensen, 2019). For this purpose, different methodologies have been explored, for example refining bonded and non-bonded potentials (Denning et al., 2011; Robertson et al., 2019; Tan et al., 2018) and adding external hydrogen terms (Kuhrova et al., 2019). The improved force fields have led to successful implementations of MD simulations on some RNA-containing systems, including conformational dynamics of folded RNAs (Banas et al., 2010; Krepl, Clery, Blatter, Allain, & Sponer, 2016; Kuhrova et al., 2016; Musiani et al., 2014; Salmon, Bascom, Andricioaei, & Al-Hashimi, 2013; Templeton & Eber, 2018; Y. Xu, MacKerell, & Nilsson, 2016) and biological catalysis (Banas, Jurecka, Walter, Sponer, & Otyepka, 2009; K. Nam, Gao, & York, 2008; K. H. Nam, Gaot, & York, 2008). Concerning the second issue, naturally occurring modified ribonucleotides parameters have now been developed and are consistent with commonly used force fields (Aduri et al., 2007; Y. Xu, Vanommeslaeghe, et al., 2016). The last issue may be most critical for the successful simulation of a protein-RNA complex. For balancing protein-RNA interactions, one strategy is to modify pair-specific corrections to non-bonded interactions by targeting related osmotic pressure (Y. Luo & Roux, 2010; J. Yoo & Aksimentiev, 2016, 2018; J. J. Yoo & Aksimentiev, 2012). Besides, polarizable force fields may capture essential physical effects for protein-RNA interactions, but their developments are not fully mature yet (Lemkul & MacKerell, 2018; Yuan, Zhang, Mills, Hu, & Zhang, 2018; C. S. Zhang et al., 2018).

Here, we use YTHDC1 protein as a model system to investigate the binding of the preferred RNA motif to m^6^A-reader proteins by protein crystallography, isothermal titration calorimetry (ITC), and explicit solvent MD simulations with a force field suitable for protein-RNA complexes. The role of upstream and downstream m^6^A-flanking nucleotides in the binding event is analyzed at the atomistic level. The MD simulations of the bound states reveal the flexibility of the 5’ terminal nucleotides, and the role in stabilizing cation-π-π interaction by the 3’ terminal nucleotides. From the unbound-state simulations, we found that the methyl group of m^6^A and 5’ terminus shift the nucleotides’ conformations into the bound-like one observed in the structures of the complex. Taken together, the crystal structures and simulation results of bound and free oligoribonucleotides provide an atomistic explanation for the roles of m^6^A-flanking ribonucleotides in the molecular recognition.

## Materials and methods

### Protein purification

The plasmid expressing the N-terminally hexahistidine-tagged YTH domain (residues 345-509) of human YTHDC1 protein was obtained as a gift from Cheryl Arrowsmith (Addgene ID: 64652). The recombinant protein was purified to homogeneity in two chromatographic steps. The protein was overexpressed for 16 hours at 24°C in Escherichia coli BL21 (DE3) cells upon induction with 0.4 mM IPTG. The cells were harvested and resuspended in the lysis buffer containing 100 mM Tris-HCl at pH 8.0, 500 mM NaCl and 10 mM imidazole. The cells were lysed by sonication and the cell lysate was clarified by centrifugation at 48,000 g for one hour and loaded onto Ni-NTA affinity column (5 mL HisTrap FF from GE Healthcare). After extensive washing with the wash buffer containing 100 mM Tris-HCl at pH 8.0, 500 mM NaCl and 50 mM imidazole the target protein was eluted with elution buffer containing 100 mM Tris-HCl at pH 8.0, 500 mM NaCl and 250 mM imidazole. The N-terminal hexahistidinetag was removed by cleavage with tobacco etch virus (TEV) protease at 1:50 ratio. The excess imidazole was removed by overnight dialysis and the sample was subjected to secondary subtractive Ni-NTA affinity chromatography step to remove the protease and uncleaved protein. Finally, the protein was subjected to a gel filtration step using Superdex 75 16/60 column in a buffer containing 10 mM Tris-HCl at pH 7.5, 150 mM NaCl and 1 mM DTT. The protein was concentrated to 10 mg/mL, flash-frozen in liquid nitrogen and stored at −80 °C for future experiments.

### Crystallography

The crystals of YTHDC1 YTH domain were obtained by mixing 1 μL protein solution at 10 mg/mL with mother liquor containing 0.1 M Bis-Tris at pH 6.5, 0.2 M ammonium sulfate and 25% PEG 3350 at 22°C in a hanging drop vapor diffusion setup. To obtain crystals of protein complexed with oligoribonucleotides (custom-synthesized and HPLC-purified by Dharmacon), the crystals were transferred to a 1 μL drop containing 10 mM oligoribonucleotide directly dissolved in 0.1 M Bis-Tris at pH 6.5, 0.2 M ammonium sulfate and 30% PEG 3350, soaked overnight at 22 °C, harvested and frozen in liquid nitrogen without additional cryoprotection.

Diffraction data were collected at the Swiss Light Source (Villigen, Switzerland) using the beamline X06DA (PXIII), and processed using XDS (Kabsch, 2010). The structures were solved by molecular replacement using Phaser program (Mccoy et al., 2007) from the Phenix package (Afonine et al., 2012). The unliganded structure of YTHDC1 (PDB ID: 4R3H) was used as a search model. The model building and refinements were performed using COOT (Emsley, Lohkamp, Scott, & Cowtan, 2010) and phenix.refine (Afonine et al., 2012). Data collection and refinement statistics are summarized in Table S1.

### Coordinates

The atomic coordinates and structure factors for the four determined complexes have been deposited in the Protein Data Bank (PBD) with the accession codes 6RT4, 6RT5, 6RT6 and 6RT7.

### Binding assays

Isothermal titration calorimetry (ITC) experiments were carried out at 18°C using MicroCal ITC200 (GE Healthcare). For the ITC experiments, the final gel filtration step of the purification of YTH domain of YTHDC1 was performed in the ITC buffer (20 mM HEPES, pH 7.5, 150 mM NaCl). Oligoribonucleotides were directly dissolved at 500-800 μM in the ITC buffer and titrated into the sample cell containing the protein at 50-60 μM. After an initial injection of 0.4 μL, 19 injections of 2.0 μL each were performed. The raw data were integrated, normalized for concentration, and analyzed using a single binding site model, provided in the MicroCal Origin software package.

### Molecular dynamics simulations

All the systems for MD simulations in this study were constructed based on the structure of the YTH domain of YTHDC1 complexed with modified pentaribonucleotide reported previously by others (PDB ID: 4R3I) (C. Xu et al., 2014). The protein YTHDC1, the RNA oligomer GG(m^6^A)CU, and crystal water molecules were kept as in the initial structure. Based on this structure, six different systems were generated, i.e., three complex systems and three structures of unbound modified oligoribonucleotides. For the three complex systems, the protein coordinates were kept unchanged, while the oligoribonucleotides were truncated to obtain the tetrameric G(m^6^A)CU and GG(m^6^A)C from the pentameric GG(m^6^A)CU, whose coordinates were available in the crystal structure. The hydrogen atoms were added by the CHARMM program (Brooks et al., 2009) and the protonation states were determined at neutral pH conditions. The complex systems were solvated in a 67 Å of rhombic dodecahedron (RHDO) TIP3P water box (Jorgensen, Chandrasekhar, Madura, Impey, & Klein, 1983) to ensure 10 Å buffer space between the macromolecular atoms and the boundary of the water box. To neutralize the system and mimic the physiological conditions, Na+ and Cl− ions at 0.15 M concentration were added to the solvated systems. Similarly, the three free RNA ligands were solvated in a 47 Å of RHDO water box with 0.15 M of NaCl. To explore the regulatory role of the m^6^A’s methyl group in RNA’s conformations, the unmethylated pentamer GGACU was also simulated in its free state.

Each simulation system was initially minimized for 10,000 steps under a series of restraints and constraints on the solute atoms to release its bad contacts and poor geometry. The minimized structure was heated to 300 K and equilibrated in NVT condition (constant volume and temperature). Finally, the structure was further equilibrated in NPT condition (constant pressure and temperature). All the equilibration phases lasted for 1 nanosecond (ns) using the CHARMM program (version 42b2) (Brooks et al., 2009). Production runs of 500 ns each were carried out in NPT conditions using the NAMD program (version 2.12) (Phillips et al., 2005). The pressure was controlled by Nosé-Hoover Langevin piston method with 200 picosecond (ps) piston period and 100 ps piston decay time (Feller, Zhang, Pastor, & Brooks, 1995; Martyna, Tobias, & Klein, 1994). The temperature was maintained at 300 K using the Langevin thermostat with a 5 ps friction coefficient. The integration time step was set to 2 fs by constraining all the bonds involving hydrogen atoms by the SHAKE algorithm. Van der Waals energies were calculated using a switching function with a switching distance from 9 to 11 Å (Steinbach & Brooks, 1994), and electrostatic interactions were evaluated using the particle mesh Ewald summation (PME) method (Essmann et al., 1995).

The CHARMM36 force field was used for protein (Huang & MacKerell, 2013) and RNA (Denning et al., 2011) molecules. For the parameterization of m^6^A, the force field for naturally occurring modified ribonucleotides was used (Y. Xu, Vanommeslaeghe, et al., 2016). Five independent runs with random initial velocities were carried out for each system for a total of 2.5 microseconds (μs). MD snapshots were saved every 20 ps along the MD trajectories for further analysis. Geometric measurements, e.g., distance, dihedrals, root mean square deviation (RMSD) and contact map analysis, were performed with CHARMM routines. All statistical figures were plotted by MATLAB (Version 2018a MathWorks, Inc.) and structural figures were generated with the PyMOL graphic software (Version 2.2 Schrödinger, LLC.).

## Results

### Flexible binding of the G_−2_G_−1_ segment of the GG(m6A)CU ligand to the YTH domain

The crystal structure of the complex between the pentamer GG(m^6^A)CU and YTHDC1 YTH domain (PDB ID: 4R3I) shows that m^6^A is fully buried in an aromatic cage consisting of Trp377 and Trp428 (Figure 1) (C. Xu et al., 2014). On the other hand, the contribution to binding of the non-modified nucleotides is less clear. The two guanosines of GG(m^6^A)CU are visible in the structure 4R3I, interacting with the surface residues of YTHDC1 (Figure 1A). The guanine bases of G−1 and G−2 form hydrogen bonds (HB) with the backbone of Val382 and Asp476, respectively (Figure 2A). Furthermore, there is a HB between the two guanines (C. Xu et al., 2014). To test the stability of these interactions, we conducted MD simulations on the complex system of YTHDC1 with GG(m^6^A)CU. The interaction analysis shows that the HB interactions are not well maintained during the simulations with large deviation from the distances in the crystal structure (Figure S1).

**Figure 1.**
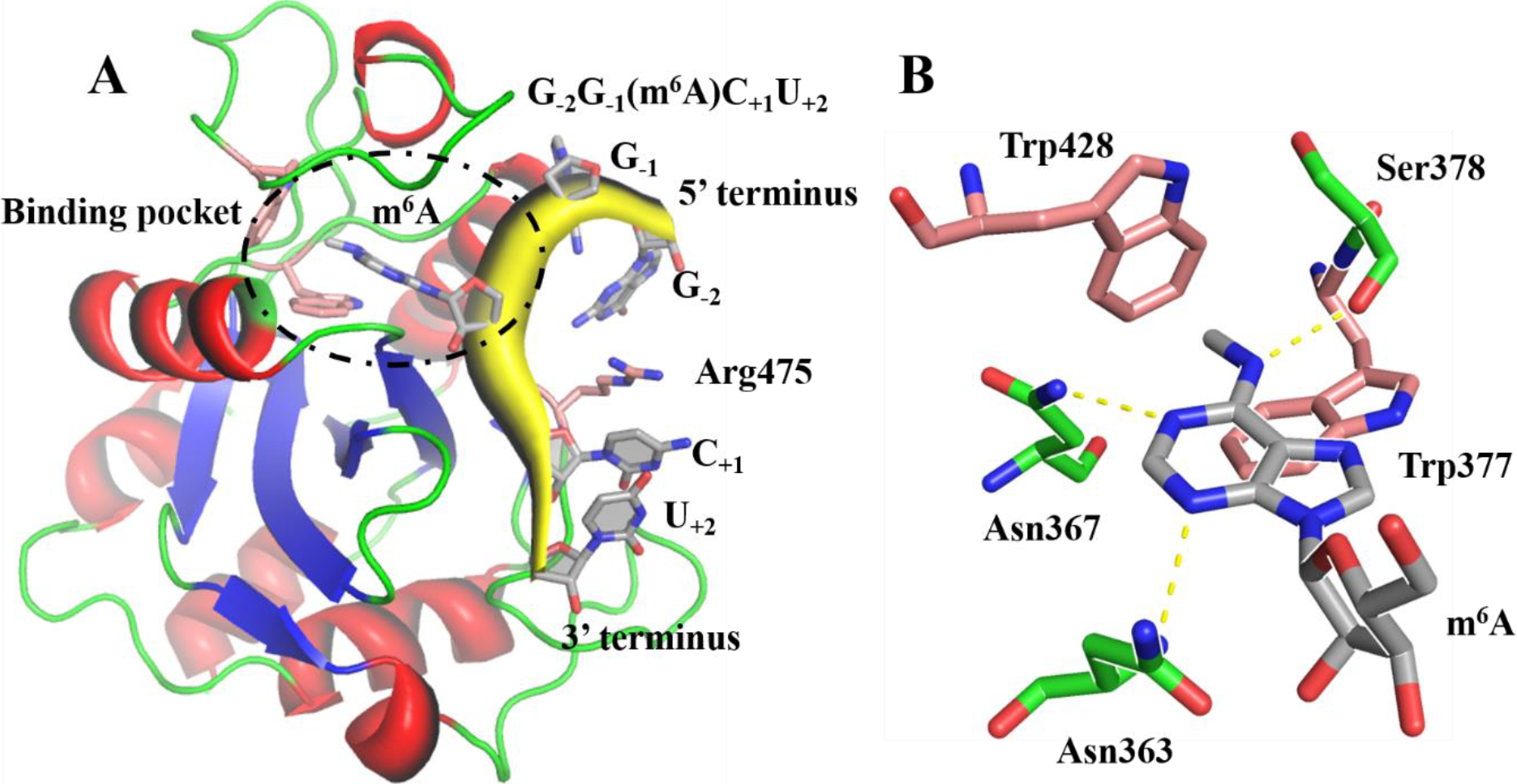
Binding mode of the RNA motif GG(m^6^A)CU to the YTHDC1 reader domain (PDB ID: 4R3I). (A) The backbones of RNA (yellow) and YTHDC1 (red, blue, and green for α-helices, β-strands, and loops, respectively) are shown in cartoon and the two tryptophanes of the aromatic cage are shown in sticks. The binding pocket of m^6^A is highlighted (black dashed circle). (B) Detailed interactions between m^6^A (carbon atoms in gray) and the pocket of YTHDC1 (tryptophane and polar residues in salmon and green, respectively). The importance of the two tryptophanes for the binding has been confirmed by site-directed mutagenesis experiments (C. Xu et al., 2014).

**Figure 2.**
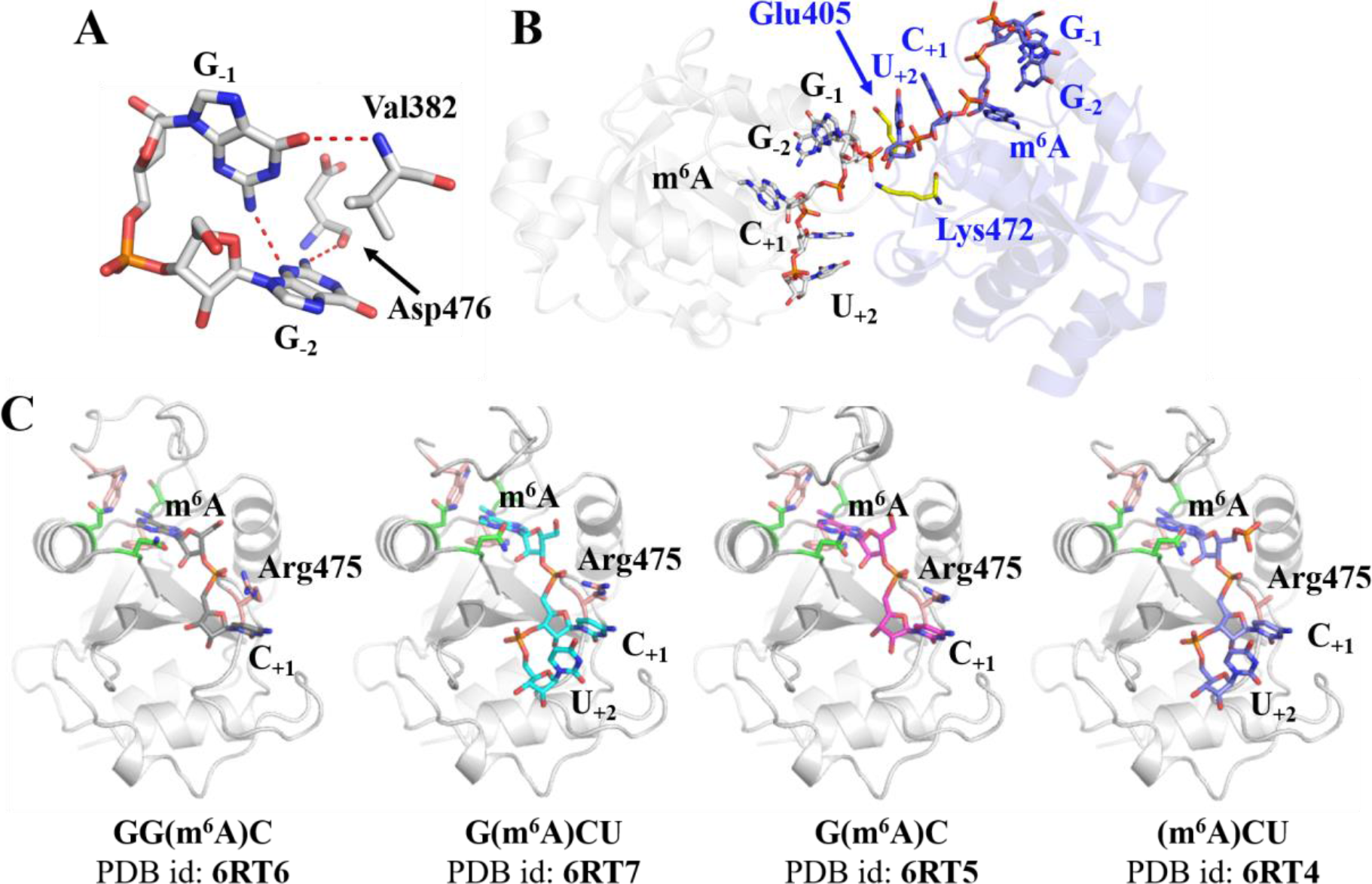
Interactions stabilized by crystal packing. **(A)** Hydrogen bonds between nucleotides (G_−1_ and G_−2_) and YTHDC1. The interaction mode was extracted from the structure 4R3I. The plausible hydrogen bonds are indicated in red dotted lines. **(B)** Two m^6^A-containing RNA oligomer-YTHDC1 complexes related by crystallographic symmetry, shown and labeled in gray and blue for clarity. Their RNA structures are presented in sticks. The protein residues that contribute the crystal packing interactions are colored in yellow. **(C)** New complex structures solved in the present study. The proteins of the complex structures are shown in gray cartoon and the bound RNAs are shown in sticks with different colors. The residues in the m^6^A binding site are shown in sticks (same colors as in Figure 1B).

We checked whether crystal packing plays a role in stabilizing the HB interactions between the nucleotides G_−1_G_−2_ and YTHDC1. In particular, several interactions are found at the interface between molecules related by crystallographic symmetry (Figure 2B). For example, towards the RNA’s 5’ end, the sugar group of G_−2_ interacts with the side chain of Glu405 of the monomer related by translation and the 2_1_ screw axis. Similarly, the phosphate linker of G_−1_ in the gray monomer interacts with the sugar group of U_+2_ in the blue monomer and is further stabilized by a salt bridge contributed by Lys472. Such contacts of oligoribonucleotides from two YTHDC1-RNA complexes are unlikely populated in aqueous environment, and therefore they are probably a consequence of crystal packing. Similar packing modes are also found in other structures of reader proteins (Figure S2).

We hypothesized that truncated variants of GG(m^6^A)CU, i.e., without terminal nucleotides, would not permit the crystal contacts mentioned above. To test this, four different RNAs were synthesized for crystallographic study, i.e., GG(m^6^A)C, G(m^6^A)CU, G(m^6^A)C, and (m^6^A)CU. We solved the crystal structures of the YTH domain of YTHDC1 in the complex with these four RNA oligonucleotides at resolutions of 1.5, 1.7, 2.3 and 1.5 Å, respectively (Table S1). The new structures mostly agree well with the structure of the complex with the pentamer RNA ligand (Figure 2C). However, the coordinates of the two guanine nucleotides, including bases, sugars and phosphates, at positions −2 and −1 with respect to m^6^A cannot be modelled in our structures due to missing electron density. By contrast, the cytosine and uridine at positions +1 and +2, respectively, show well defined electron density. Thus, our co-crystal structures suggest that the two guanine ribonucleotides upstream of m^6^A do not form stable interactions with the protein, while the downstream motif CU forms conserved interactions with the protein. These findings are consistent with our hypothesis on the stabilization of the untruncated GG(m^6^A)CU due to crystal packing in the YTHDC1 structure (PDB id: 4R3I).

### 5’ and 3’ terminal nucleotides contribute to the affinity of GG(m^6^A)CU to YTHDC1

The four crystal structures have provided evidence that the G_−2_G_−1_ dinucleotide segment does not form stable interactions with YTHDC1 while the segment C_+1_U_+2_ shows conserved contacts with the protein. However, it is not clear if and how much these terminal nucleotides contribute to the binding of GG(m^6^A)CU to YTHDC1. To address this question, we measured dissociation constant (K_d_) values of three representative oligomers, viz., GG(m^6^A)CU, G(m^6^A)CU, and GG(m^6^A)C using isothermal titration calorimetry (ITC) (Figure 3 and Table S1). We obtained a K_d_ value of 0.5 μM for GG(m^6^A)CU, which is similar to a value reported previously (2.0 μM) (C. Xu et al., 2014). The U_+2_-deficient tetramer, i.e., GG(m^6^A)C, is less potent by a factor of about 20 than the pentamer GG(m^6^A)CU. Such binding reduction can be intuitively understood from the crystal structures, namely U_+2_ forms π-π stacking with C_+1_ and thus stabilizes interactions of the 3’ end with YTHDC1. Surprisingly, the G_−2_-deficient oligomer, i.e., G(m^6^A)CU, shows significant reduction of affinity (∼7 times compared to the pentamer) although G_−2_ does not form stable interactions with YTHDC1 in our crystal structures and MD simulations (Figure 1C and S1). Thus, further investigation is needed to understand how G_−2_ impacts the binding affinity (See below).

**Figure 3.**
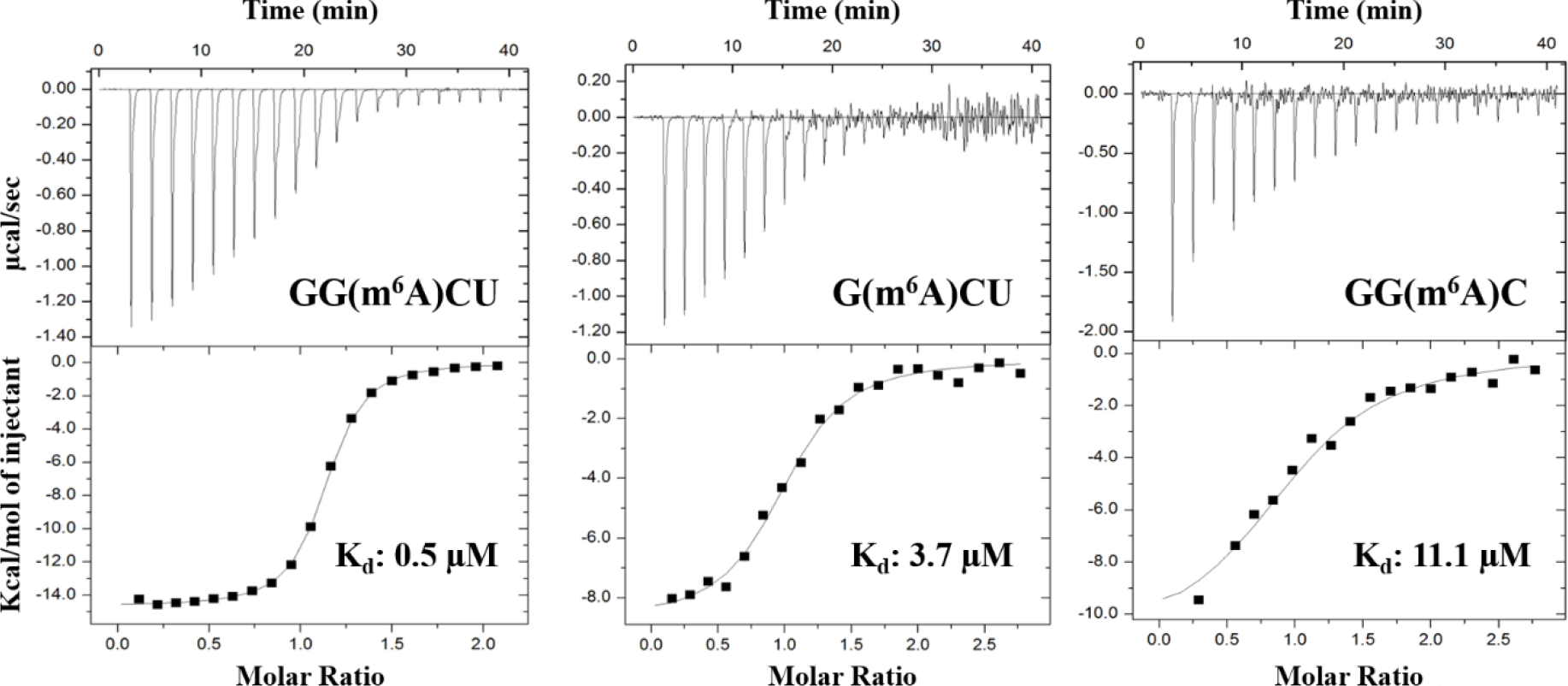
Isothermal titration calorimetry measurements of the binding of the RNA oligomers GG(m^6^A)CU, G(m^6^A)CU, and GG(m^6^A)C to YTHDC1. The raw ITC curves and binding isotherms with the best fit are shown in the top and bottom panels, respectively. The truncation of one nucleotide at either 5’- or 3’-terminus of GG(m6A)CU deteriorates the affinity.

### MD simulations of liganded YTHDC1 reveal flexibility of the nucleotides upstream of m^6^A

Three complexes with ligand GG(m^6^A)CU, G(m^6^A)CU, and GG(m^6^A)C were investigated by MD simulations to analyze their relative flexibility. In all simulations, the two guanine nucleotides upstream of m^6^A show large deviations compared to their crystal poses (Figure 4 and Figure S3). This result is consistent with the lack of electron density for this RNA segment. During the MD simulations, the two guanine nucleotides do not show stable and specific interactions with YTHDC1 in contrast with the crystal structure (PDB code 4R3I) (Figure 2A). Instead, the two nucleotides are dynamically coupled for a long time by the π-π interaction between their bases. This is illustrated by the contact map analysis, in which G_−1_ and G_−2_ interact with each other in approximately half of the samplings (lower triangles of Figure 5).

**Figure 4.**
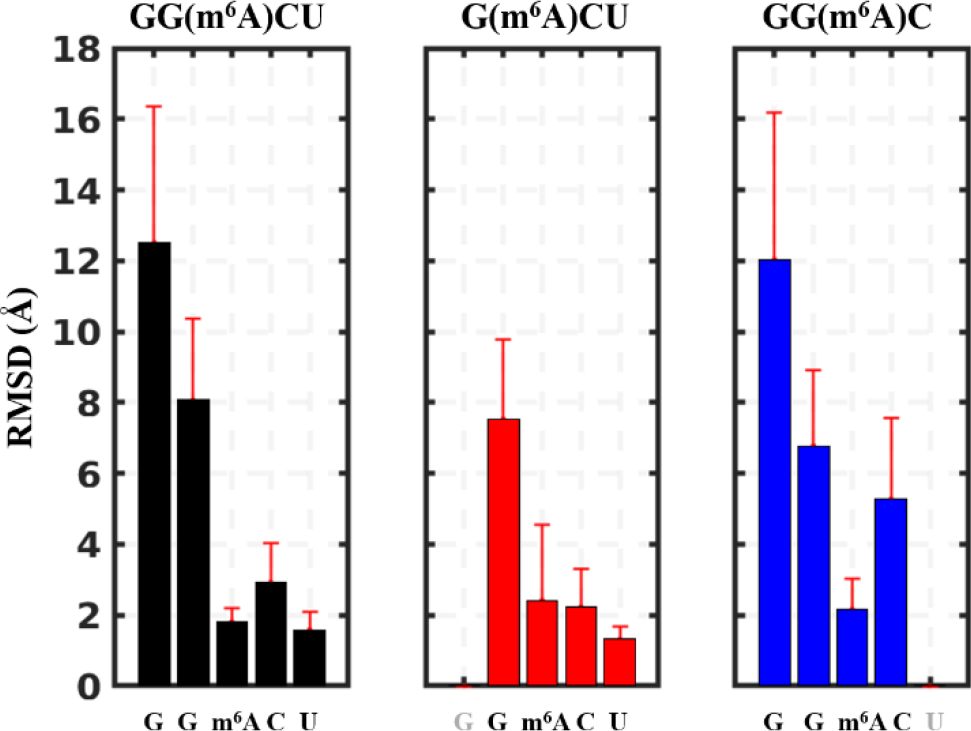
RMSD profile of RNA oligomers in the bound state. The plots show average RMSD value and standard deviation calculated over the cumulative sampling of five trajectories for each simulation system. To calculate the RMSD value, heavy atoms of nucleotides in their crystal structure (PDB ID: 4R3I) were used as reference. All snapshots from MD simulations were superimposed to the protein backbone of the crystal structure excluding C- and N-terminal segments.

**Figure 5.**
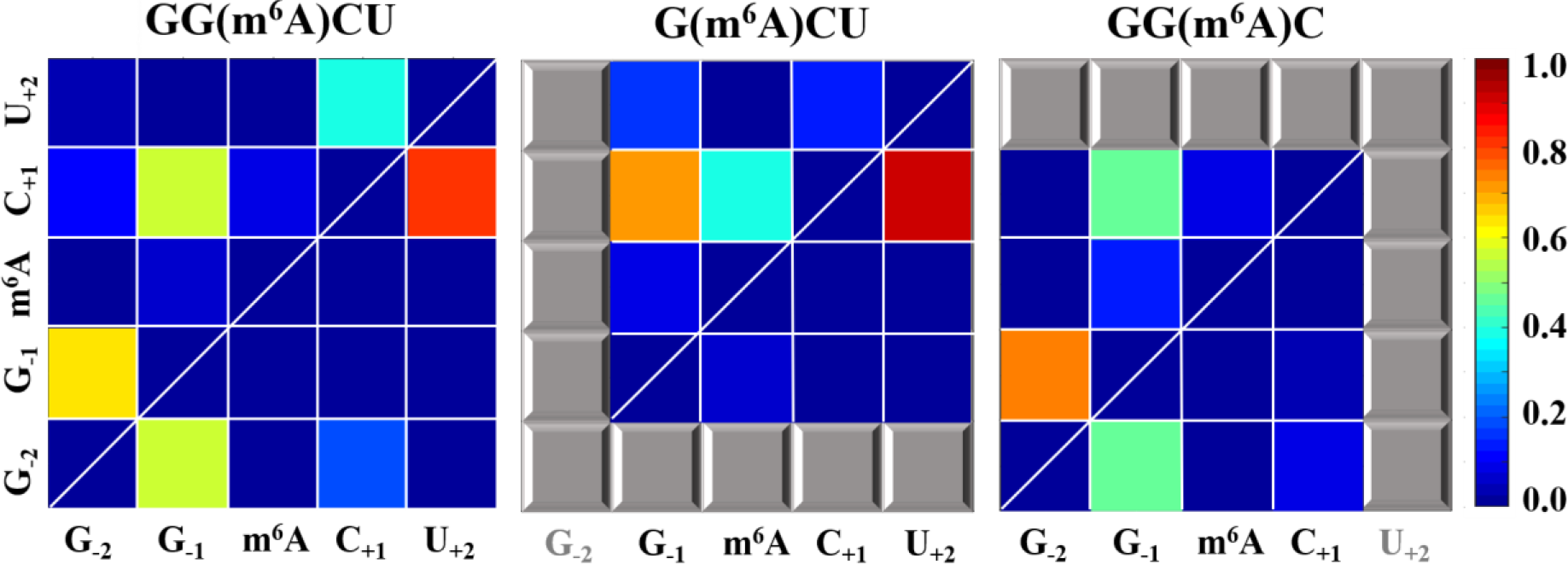
Contact maps of m^6^A-containing RNA oligomers in their free state (upper, left triangle) and in the complex with YTHDC1 (lower, right triangle). Pairwise contact frequencies are evaluated by the distance of center of mass between two bases and a cutoff value of 5 Å. The color bar represents the frequency of contact between two nucleotides, ranging from dark blue (no interaction) to dark red (persistent interaction). The nucleotide pairs which are not included in the systems are presented in gray squares.

The tetraribonucleotide ligands allow us to assess the role of the terminal G_−2_ and U_+2_, without which m^6^A-containing RNAs show reduced binding affinity compared to the pentamer. Although the nucleotide G_−2_ was flexible during the simulations of the complex, it helps stabilizing the pose of m^6^A in the binding pocket of YTHDC1. Without the terminal guanine nucleotide at position −2, m^6^A experiences slightly larger deviations and fluctuations than in G_−2_-containing ligands (Figure 4 and Figure S4). The larger RMSD values are due to the partial dissociation of m^6^A from the YTHDC1 binding pocket in two of the five trajectories of G(m^6^A)CU, which can be monitored by three key hydrogen bonds between m^6^A and YTHDC1 (Figure S4). Noteworthy, if the dissociated snapshots are neglected in the system G(m^6^A)CU, the RMSD value of m^6^A is nearly unchanged compared to other two systems (Figure S5). Therefore, the reduced affinity of the G_−2_ deficient ligand may not be plainly explained based on simulation results of the bound states. Other factors causing the affinity reduction will be investigated in the later sections.

Concerning the nucleotides downstream of m^6^A, our crystal structures and MD simulations provide evidence that C_+1_ and U_+2_ form more stable interactions with YTHDC1 than G_−1_ and G_−2_. The downstream C_+1_ establishes cation-π and π-π interactions with Arg475 and U_+2_, respectively. A previous study has confirmed the importance of the cation-π interaction for the molecular recognition between YTHDC1 and GG(m^6^A)CU by mutating Arg475 to phenylalanine or alanine with 10-fold and 100-fold reduction, respectively, compared to the wild type (C. Xu et al., 2014). However, the role of U_+2_ has not be confirmed as it is stabilized by the crystal packing in the structure 4R3I (Figure 2B). The simulations for the tetrameric ligand GG(m^6^A)C show that the C_+1_-Arg475 interaction becomes less stable than in the case of the ligand with U_+2_. Consistently, the RMSD of C_+1_ becomes much larger and fluctuates more than that of the U_+2_-containing oligomers (blue bars in Figure 4). This observation indicates that U_+2_ indeed stabilizes the interaction between C_+1_ and Arg475. As shown in Figure S6, the contact distance for C_+1_ and Arg475 is locally distributed at 3.8 Å for U_+2_-containing ligands (black and red lines in the Figure S6), while it becomes loosely distributed in the U_+2_-deficient ligand (the blue line). These structural and dynamic analyses rationalize the role of U_+2_ and the reduced binding of the tetrameric ligand GG(m^6^A)C (shown by ITC data).

The C_+1_U_+2_ segment specifically interact with a positively charged shallow pocket on the protein surface and may play comparably important role in the protein-RNA recognition compared to m^6^A. An equivalent mutation to Arg475Ala was also reported for a homologous protein YTHDF2, i.e., Arg527Ala. This mutation showed approximately 25-fold reduction of the binding affinity for its RNA oligoribonucleotides compared to the wild type (Zhu et al., 2014). Moreover, the C_+1_U_+2_ segment binds to a well-defined pocket on the YTHDC1 surface, consisting of four positively charged residues, i.e., Lys361, Arg404, Lys472, and Arg475 (Figure 1A). The binding data and structural analysis suggest that the interaction between the C_+1_U_+2_ segment and the reader protein is specific rather than an undirected charge-charge interaction. In addition, Zhu *et al.* suggested that a two-step model fits the data from a fluorescence polarization assay better than the one-step model for the binding of YTHDF2 to an m^6^A-containing oligoribonucleotide (Zhu et al., 2014). They proposed that the binding of m^6^A-flanking nucleotides facilitates the recognition of m^6^A. Integrating this two-step model with our results from ITC and MD simulations, we infer that the C_+1_U_+2_ segment play an essential role in the association process of GG(m^6^A)CU to DC1 comparable to that of m^6^A. However, it is still not clear which the first motif is to be recognized by the reader protein, m^6^A or C_+1_U_+2_ segment, and this question requires further studies.

### Conformational heterogeneity of m^6^A-containing RNAs in bound and free states

To rationalize relative binding affinities, we analyzed conformations of three RNA oligomers in their bound and free states. Significant differences are found in the intramolecular contact pattern between the bound and free states (Figure 5). The bound RNAs show reduced contacts between nucleotides compared to their free states. For example, three pairs of nucleotides in GG(m^6^A)CU’s free state are in contact for more than 40% of all snapshots. Its bound state, by contrast, loses one stable interaction pair, i.e., G_−1_-C_+1_ interaction pair (Figure 5). That is because the centrally located m^6^A of the bound state firmly anchors in the aromatic cage of YTHDC1. Thus, the pentameric GG(m^6^A)CU is divided into two independent segments, i.e., G_−1_-G_−2_ and C_+1_-U_+2_, separated by m^6^A. Two nucleotides belonging to the same segment are frequently coupled, but nucleotides from different segments interact much less (lower triangles in Figure 5).

The purine base of m^6^A fully interacts with the aromatic cage of YTHDC1. Therefore, it is reasonable to expect that m^6^A should be sufficiently exposed to the solvent before the association with the protein, so that its binding pocket can effectively recognize m^6^A. The contact maps show that m^6^A in the G_−2_-containing ligands, i.e., GG(m^6^A)CU and GG(m^6^A)C, is not in frequent contact with other nucleotides in the unbound state. By contrast, m^6^A in G(m^6^A)CU does interact with C_+1_ (upper triangles in Figure 5). The distribution of m^6^A-C_+1_ distance for G(m^6^A)CU is more populated for values around 5 Å than for the other two ligands (Figure S7).

To detect how the exposure of m^6^A varies in different ligands, we measured the solvent accessible surface area (SASA) of m^6^A in the free state (Figure S8A). The two G_−2_-containing ligands show a similar distribution of SASA, which is mostly populated around 180 Å^2^ with a second peak at 200 Å^2^. By contrast, G(m^6^A)CU populates its main peak at 150 Å^2^. Such difference in SASA is caused by the frequent contact between m^6^A and C_+1_ (the upper triangle for G(m^6^A)CU in Figure 5). Taken together, the G_−2_ nucleotide plays a role in the exposure of m^6^A from the RNAs in the unbound state, and thereby benefits the recognition of m^6^A by its reader proteins. To study if the methyl group of m^6^A facilitates the base exposure, we also compared the oligomers GG(m^6^A)CU and GGACU in their unbound states. Again, a significant difference was found in the distribution of the SASA, with the unmethylated oligomer showing a less exposed adenine than its N6-methylated counterpart (Figure S8B).

### Conformational free-energy profiles of m^6^A-containing RNAs in the unbound state

Multiple MD simulations of the unbound state of GG(m^6^A)CU, G(m^6^A)CU, and GG(m^6^A)C were carried out to analyze potential differences in the most populated conformers. The simulations were used to construct free-energy profiles with the RMSD of (m^6^A)C relative to its crystal pose (Changeux & Edelstein, 2011) as the progress variable. The free-energy profiles show that oligomers in aqueous solution adopt multiple meta-stable conformations, which appear as local minima of the free-energy profiles (Figure 6). The ligands GG(m^6^A)CU and GG(m^6^A)C show a similar conformational propensity. There is approximately a 1 kcal/mol free-energy difference between their global minimum and the bound-like local minimum (black and blue lines in Figure 6). By contrast, G(m^6^A)CU experiences a ∼2 kcal/mol of energy penalty from the unbound to the bound state. The conformational free-energy difference indicates that the former two ligands have less energy penalty for redistributing their conformations to the bound-state than G(m^6^A)CU. It is useful to introduce a two-state approximation in which all conformations of an RNA oligomer are divided into either bound or unbound states. The two-state model shows a significant energy difference between G_−2_-containing oligomers and G(m^6^A)CU irrespective of the RMSD threshold (Figure S9). A similar difference was also found by comparing the PMF profiles of unbound GG(m^6^A)CU and GGACU (Figure S8C), indicating that the methyl group of m^6^A helps in adopting the bound conformation.

**Figure 6.**
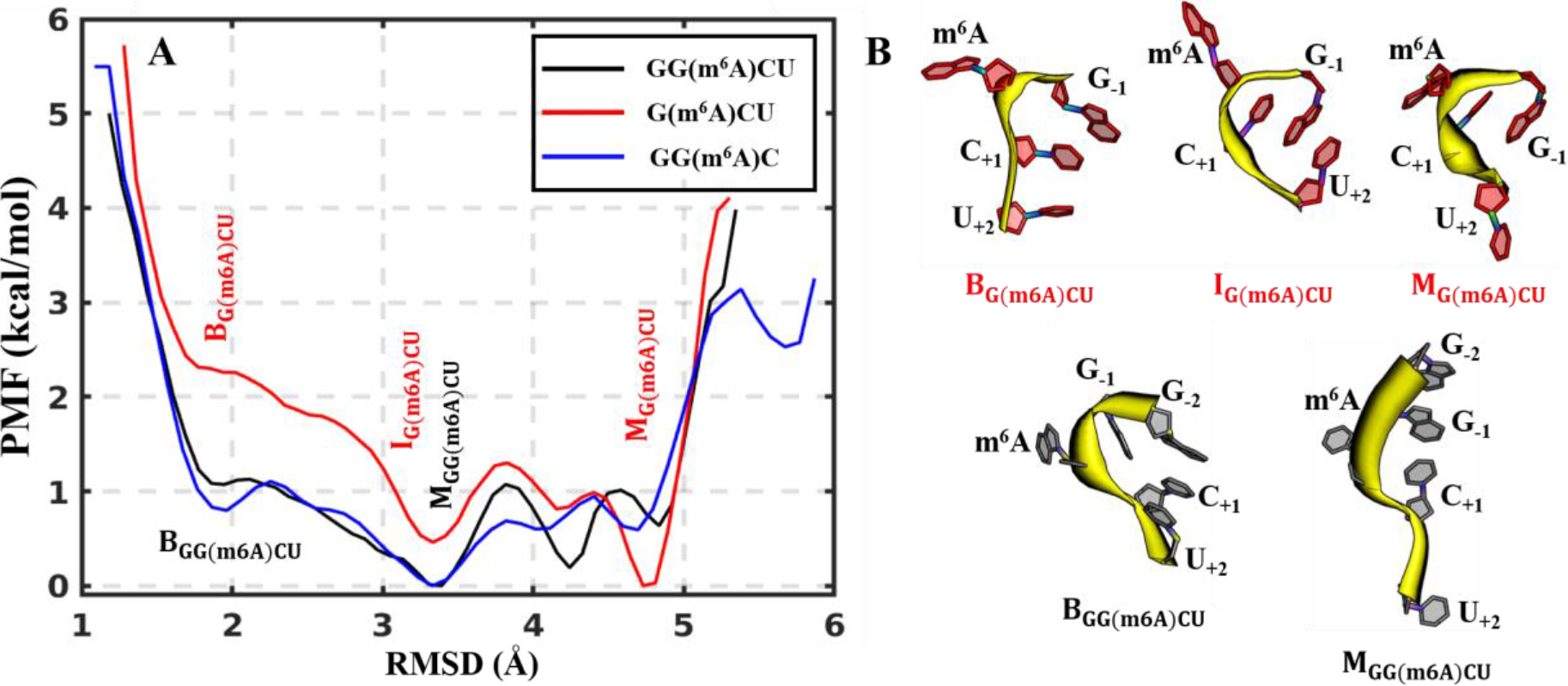
Conformational landscape of m^6^A-containing RNA oligomers in the unboun d state. (A) Potential of mean force (PMF) along the RMSD from the bound state. The structure of the RNA segment present in the three simulated oligonucleotides, i.e., (m^6^A)C, in the complex with YTHDC1 (PDB code 4R3I) is selected as reference for the bound state. The PMF is calculated as 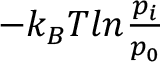, where *k*_*B*_ is Boltzmann factor, *T* is temperature (300 K), *p*_*i*_ is the population in bin *i*, and *p*_0_ is the probability of the most populated bin (with a 0.1 Å of bin width). The labels M, I, and B indicate mostly populated, intermediate, and bound-like conformational states, respectively. (B) Representative structures of different conformational states of free G(m^6^A)CU (top) and GG(m^6^A)CU (bottom).

The analysis on meta-stable conformations suggests an important role of G_−2_ in maintaining the bound-like conformations and exposing m^6^A to the bulk water. For example, for the oligomer GG(m^6^A)CU, the global minimum conformation along the free-energy profile not only maintains a bound-like pose but also exposes m^6^A to the solvent (M_GG(m6A)CU_ in Figure 6B). In this conformation, G_−2_ interacts with G_−1_ in a π-π interaction and thereby frees m^6^A to expose to the bulk water. In the most stable conformation of G(m^6^A)CU, G_−1_, m^6^A and C_+1_ stack together and thus m^6^A is less exposed to the solvent (the upper panel in Figure 6B). Although the conformational change from M_G(m6A)CU_ to I_G(m6A)CU_ increases the solvent exposure of m^6^A, C_+1_ and G_−1_ become packed with each other, resulting in a detrimental pose for its binding to YTHDC1. Finally, to transform this intermediate state to the bound-like one, the ligand G(m^6^A)CU has to pay a higher free-energy cost than GG(m^6^A)CU. Thus, G_−2_ is a key nucleotide for modulating the conformational preference of m^6^A-containing RNAs in the unbound state.

## Discussion

We aimed to elucidate the roles of m^6^A-flanking nucleotides in the molecular recognition of YTHDC1 to its RNA motif GG(m^6^A)CU by atomistic simulations, X-ray crystallography, and isothermal titration calorimetry. The selected model oligoribonucleotide is biologically relevant because the sequence is preferentially recognized by all five human m^6^A reader proteins (A. Li et al., 2017; F. D. Li et al., 2014; Shi et al., 2017; C. Xu et al., 2014). To investigate the flexibility of terminal nucleotides, the structures of the YTH domain of YTHDC1 in the complex with GG(m^6^A)C, G(m^6^A)CU, G(m^6^A)C, and (m^6^A)CU were solved at high resolution. ITC binding experiments were utilized to examine the contribution of 5’ and 3’ terminal nucleotides to the affinity of GG(m^6^A)CU for YTHDC1. MD simulations were carried out for bound and unbound states of three representative RNA ligands, i.e., GG(m^6^A)CU, G(m^6^A)CU, and GG(m^6^A)C. The experimental and simulation results complement each other and provide a consistent and detailed picture of the binding mechanisms.

### Crystal packing artificially stabilizes the poses of m^6^A-flanking oligoribonucleotides bounded to reader proteins

The contacts due to crystal packing play a non-negligible role in stabilizing the binding pose of m^6^A-flanking nucleotides with YTHDC1. As shown in Figure **2**B, two 5’ terminal nucleotides (G_−2_G_−1_) in an asymmetric unit interplay with another unit by multiple artificial contacts due to crystal packing, including hydrogen bonds and salt bridges. Although some interactions are also found between YTHDC1 and the m^6^A-flanking nucleotide in the same unit (Figure 2A), they may not be strong enough to maintain the observed crystal binding pose. The crystal packing could be still a critical stabilizing factor for the crystal pose. For testing this hypothesis, we soaked varied m^6^A-containing oligoribonucleotides into apo crystals of YTHDC1. Its space group does not permit artificial intramolecular contacts of oligoribonucleotides from different asymmetric units but still leave enough space for their natural interactions with the protein in the same unit. As a result, the G_−2_G_−1_ segment is disordered in all soaking structures, indicating weak interactions between them and the protein surface. The MD simulations further support this observation as the G_−2_G_−1_ segment significantly deviates from the crystal pose and shows substantial flexibility (Figure 4 and S1). Our evidence suggests a dynamical model for the G_−2_G_−1_ segment binding to YTHDC1 against the static model which seems to emerge from the previously published structure (4R3I)(C. Xu et al., 2014).

The available structural data suggest that the binding poses of 5’ terminal nucleotides are often stabilized by crystal packing, and thus their crystal poses, and corresponding interactions should be carefully interpreted. Structural artifacts due to crystal packing are observed not only for YTHDC1, but also for other reader domains, leading to well-defined binding poses of m^6^A-flanking nucleosides, which otherwise are flexible in solution. We found in the PDB Data Bank another two m^6^A reader protein structures that are complexed with m^6^A-containing nucleotides, and they show multiple packing contacts between their m^6^A-flanking nucleotides and the protein surface (Figure S2). For example, the G_−2_G_−1_ segment in the structure 4RCJ (YTHDF1) interacts with the protein residue Ser461 and Ala462 of another asymmetric unit by multiple hydrogen bonds. Moreover, in the structure of YTH domain of MRB1 protein from Z. roux (PDB code 4U8T), the m^6^A-flanking nucleotides from different protein copies are even tangled, resulting in substantial artificial contacts (S. K. Luo & Tong, 2014). In addition, in an in-house YTHDF3 structure (to be published), the G_−2_G_−1_ segment is also stabilized by its own copies from another asymmetric unit.

### Conformational analysis of oligoribonucleotides in their unbound state is essential for understanding the recognition mechanism

Our study reveals that the 5’ terminus of GG(m^6^A)CU contributes to the affinity for YTHDC1 mainly because of the role of the 5’ terminus in the unbound state rather than its direct interactions with the protein in the bound state. The importance of the terminal nucleotide G_−2_ for the affinity is supported by our ITC measurements, which show a 7-fold reduction of the G(m^6^A)CU motif with respect to the pentameric GG(m^6^A)CU (Figure 3 and Table S2). As mentioned above, the attenuated binding affinity is not due to the missing interactions between the 5’ terminus and YTHDC1’s protein surface. It is rationalized by MD simulations of different oligoribonucleotides in aqueous solution. The corresponding free energy landscape analysis demonstrates that the 5’ terminal nucleotide G_−2_ contributes to the binding by facilitating the adoption of bound-like conformations of the RNA oligomers (Figure 6), thereby reducing the free energy expense for the association process. These results show that information from both complex structures and conformations of oligoribonucleotides in aqueous solution are indispensable to study the recognition mechanisms of reader proteins and their cognate RNA binders. This is because an oligoribonucleotide ligand has much more degrees of freedom compared to a normal-size small-molecule ligand in aqueous solution.

The conformational analysis suggests a conformational selection model. In this model, we assume that the binding pocket of YTHDC1 (i.e., the aromatic cage) is relatively rigid with only one most populated conformation. This prevalent conformation is competent for recognizing the m^6^A-containing oligoribonucleotides in its apo form. During the conformational selection by the protein, those oligoribonucleotides motifs that mainly populate the bound state in aqueous solution will pay less penalty for the conformational change than those predominating bound-resistant conformations. This model explains why GG(m^6^A)CU binds stronger than G(m^6^A)CU, thus elucidating the role of 5’ terminus G_−2_ in the binding. Our analysis also shows that m^6^A is mostly solvent exposed in the bound-like conformations (Figure S8), which is prerequisite for the recognition by the aromatic cage. In fact, the adenine also tends to be more solvent exposed in the methylated GGACU than in the unmethylated counterpart (Figure S8B). This suggests that the methyl group not only contributes the interactions of oligoribonucleotides with reader proteins but also facilities the nucleotides adopting a bound-like conformation. The capacity of the methyl group for the conformational regulation is not only limited to explain the recognition mechanism for reader proteins. The exposed conformation of methylated adenine would be also required to accommodate it in the active site of m^6^A-specific demethylase FTO (X. Zhang et al., 2019).

### Differential and shared recognition mechanisms between YTHDC1 and other reader proteins

The five human m^6^A reader proteins can be classified into two classes according to their binding pocket features, with YTHDC1 and YTHDC2 in one class and YTHDF1, YTHDF2 and YTHDF3 in the other (Figure S10). Here, we select YTHDC1 and YTHDF1 as representatives in the two classes. The first difference is in the side chain that interacts with the N1 position of m^6^A (Figure S10). Asn367 of YTHDC1 interacts with the N1 atom by donating a hydrogen atom, whilst the corresponding side chain in YTHDF1 is Asp401. The interaction between the N1 atom and the asparagine side chain is more favorable than aspartic acid (C. Xu et al., 2015). Second, the two reader proteins have different gate keeper residues at the border of their binding pockets, i.e., Asn363 and Tyr397 in YTHDC1 and YTHDF1, respectively. For the latter, the Tyr397Ala mutation abolishes its binding to GG(m^6^A)CU (C. Xu et al., 2015). The X-ray structure of holo YTHDF1 shows that Tyr397 forms π-π stacking with the 5’ terminal nucleotides (C. Xu et al., 2015). In contrast to YTHDC1, such specific interaction probably restrains the 5’ terminus dynamics on YTHDF1.

The two classes of reader proteins share two structural features, i.e., the aromatic cage for recognizing m^6^A and a positively charged shallow pocket for interactions with the 3’ terminal nucleotides. The two conserved features suggest a common mechanism for the binding of reader proteins to a m^6^A-containing RNA. First, the aromatic cage permits the selective recognition of m^6^A. The indole side chain of one of the tryptophans provides a π-π stacking with the modified adenine while the other indole is at optimal van der Waals distance with the methyl group. Our simulations show that GG(m^6^A)CU tends to expose its m^6^A to the solvent in its free state comparing to its unmethylated counterpart (Figure S8B). The m^6^A exposure provides a specific geometry for fitting the aromatic cage in all reader proteins, and thereby distinguishing unmethylated RNAs. Second, the surface pocket adjacent to the aromatic cage conservatively contains two charged residues in all five reader proteins, i.e., Arg475 and Lys361 for YTHDC1 (Figure S10). Their importance for the binding of m^6^A-containing RNAs has been verified in several reader proteins (S. K. Luo & Tong, 2014; C. Xu et al., 2015; C. Xu et al., 2014). The 3’ terminal nucleotides contribute to the protein-RNA binding by specific association to the positively charged pocket on the YTHDC1 surface. The structural features suggest a two-step binding mode, in which the electrostatically-driven association of the 3’ terminal nucleotides facilitates the further recognition and formation of optimal contacts between m^6^A and the aromatic cage (Zhu et al., 2014). Such mechanism might be common to the five human m^6^A reader proteins as they share a positive electrostatic potential on the surface that recognizes the 3’ terminus nucleotides (Figure S11). The inverse sequence of events is less probable considering that electrostatic attraction has a longer range than van der Waals interactions.

## Accession number

PDB IDs: 6RT4, 6RT5, 6RT6 and 6RT7.

## Acknowledgements

The use of beamlines and user support at Swiss Light Source are gratefully acknowledged. We thank the Swiss National Supercomputing Centre (CSCS) in Lugano for providing the computational resources. We also thank Dr. Xiang Wang for valuable discussions.

