## Supplementary files for "Flexible binding of m^6^A reader protein YTHDC1 to its preferred RNA motif"

Supplementary Table S1. Data collection and structure refinement statistics of the RNA/YTHDC1 structures.

| **PDB code** | **6RT4** | **6RT5** | **6RT6** | **6RT7** |
|---|---|---|---|---|
| **Oligoribonucleotide** | ($m^6A$)CU | G($m^6A$)C | GG($m^6A$)C | G($m^6A$)CU |
| **Data Collection:** | | | | |
| **Beamline** | SLS PXIII | SLS PXIII | SLS PXIII | SLS PXIII |
| **Space group** | $P\ 2_1$ | $P\ 2_1$ | $P\ 2_1$ | $P\ 2_1$ |
| **Cell dimensions: a, b, c (Å)** | 40.105, 103.503, 41.955 | 39.787, 103.655, 41.779 | 39.913, 103.739, 41.983 | 39.85, 103.46, 42.007 |
| **α, β, γ (°)** | 90, 104.692, 90 | 90, 104.761, 90 | 90, 104.505, 90 | 90, 104.893, 90 |
| **Resolution (Å)** | 38.87-1.49 (1.58-1.49) | 40.41-2.30 (2.44-2.30) | 44.46-1.46 (1.55-1.46) | 40.61-1.73 (1.83-1.73) |
| **Unique observations** | 53214 (8210) | 14246 (2119) | 56905 (9052) | 33768 (5348) |
| **Completeness** | 97.7 (93.9) | 97.8 (91.1) | 99.4 (97.9) | 98.2 (97.0) |
| **Redundancy** | 3.47 (3.26) | 4.48 (4.50) | 3.36 (3.27) | 3.38 (3.31) |
| **Rmerge** | 0.037 (0.597) | 0.043 (0.147) | 0.060 (1.691) | 0.09 (1.114) |
| **I/σI** | 17.15 (1.90) | 26.26 (9.86) | 12.33 (0.80) | 9.19 (1.10) |
| **Refinement** | | | | |
| **$R_{work}/R_{free}$** | 0.204/0.230 | 0.209/0.275 | 0.206/0.220 | 0.220/0.271 |
| **RMS deviations bonds (Å)** | 0.005 | 0.007 | 0.006 | 0.006 |
| **RMS deviations angles (degree)** | 0.785 | 0.911 | 0.839 | 0.821 |
| **Ramachandran Favored (%)** | 97.88 | 96.79 | 98.43 | 97.8 |
| **Ramachandran Disallowed (%)** | 0.31 | 0.31 | 0.31 | 0 |

Supplementary Table S2. Thermodynamic parameters derived from ITC measurements of different RNAs on YTHDC1.

| | GG($m^6$A)CU | G($m^6$A)CU | GG($m^6$A)C |
|---|---|---|---|
| N | 1.10 ± 0.00 | 0.99 ± 0.02 | 1.00 (fixed) |
| $K_a$ (× $10^5$ $M^{-1}$) | 21.3 ± 0.8 | 2.7 ± 0.4 | 0.9 ± 0.2 |
| $K_d$ (µM) | 0.5 ± 0.0 | 3.7 ± 0.5 | 11.1 ± 2.5 |
| ΔG (kcal/mol) | -8.5 ± 0.0 | -7.3 ± 0.1 | -6.6 ± 0.1 |
| ΔH (kcal/mol) | -14.7 ± 0.0 | -8.8 ± 0.2 | -11.1 ± 0.4 |
| -TΔS (kcal/mol) | 6.2 ± 0.0 | 1.5 ± 0.2 | 4.5 ± 0.4 |

The table summarizes the calculated binding affinity (Ka), stoichiometry (N), enthalpy (ΔH) and entropy (-TΔS). The given standard deviation is calculated from the ITC curve fitting by MicroCal Origin software. For GG($m^6$A)C, the stoichiometry was fixed to 1 because of weak binding.

Supplementary Figure S1. Interaction distance between nucleotides ($G_{-1}$ and $G_{-2}$) and YTHDC1. Plausible hydrogen bonds in the complex of YTHDC1 with GG($m^6$A)CU shown in Figure 2A are characterized by distance. The distance is measured between two heteroatoms corresponding to acceptor and donor atoms for defining a hydrogen bond. The time series of five independent runs are separated by vertical lines.

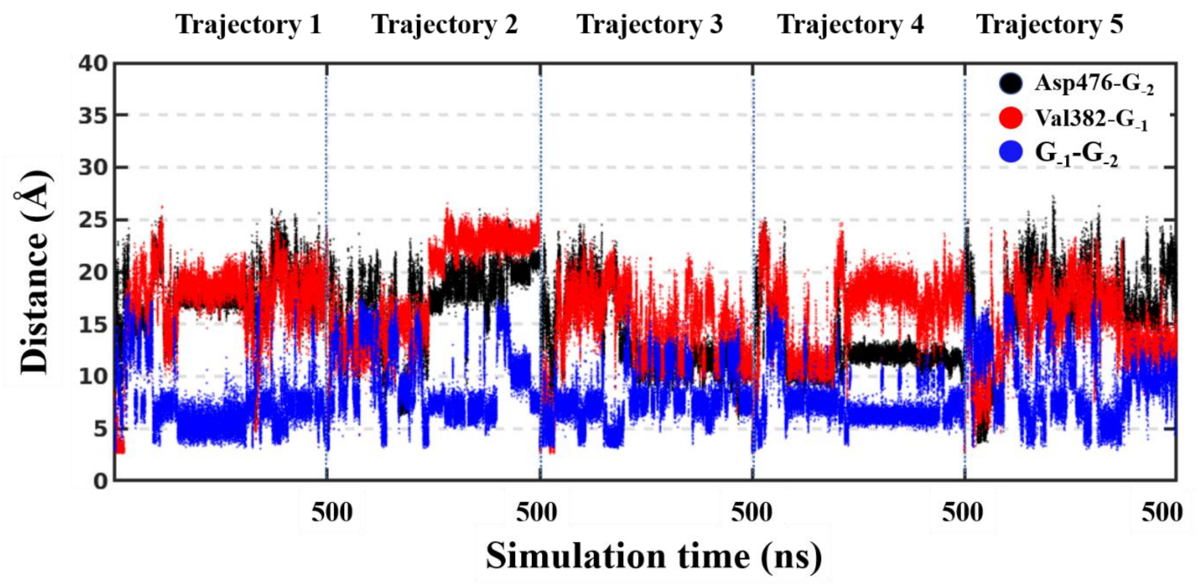

Supplementary Figure S2. Crystal packing are shown in different m$^6$A reader proteins. (A) The YTHDF1 structure (PDBid: 4RCJ). The structures from different asymmetric units are shown in green and magenta cartoon. (B) The *Z. rouxii* MRB1 structure (PDBid: 4U8T). The structures from different monomers are colored in green and magenta.

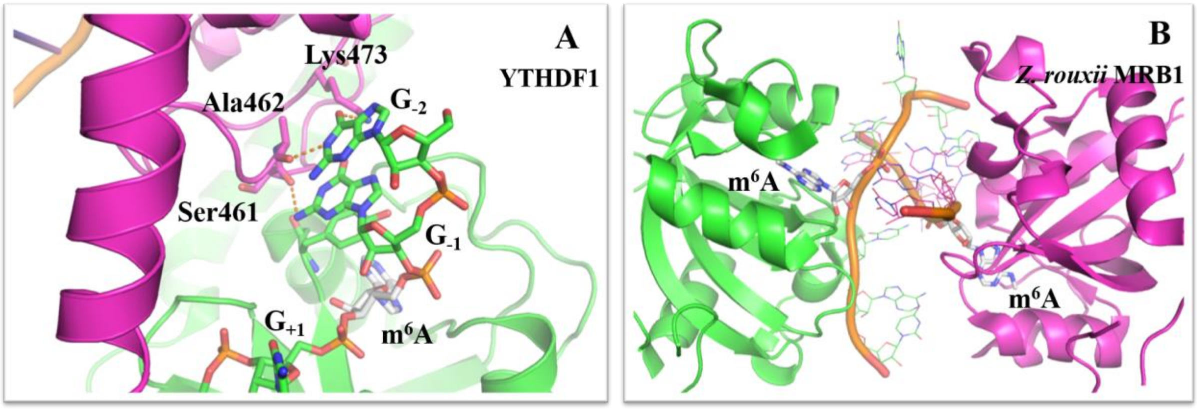

Supplementary Figure S3. Root Mean Square Fluctuation (RMSF) of RNA ligands in their bound states. The snapshots for this calculation were collected from 5 independent trajectories for each specific system. To calculate RMSF values, heavy atoms of nucleotides in their average positions (over 5 independent trajectories) were used as a reference. All snapshots of each system were superimposed to the respective crystal structure using the backbone of the protein's rigid part (by excluding C- and N-terminals), and the RMSFs were calculated based on the corresponding RNA ligands. The name of each nucleotide is written in the bottom of each sub-figure, and missing nucleotides are colored in gray.

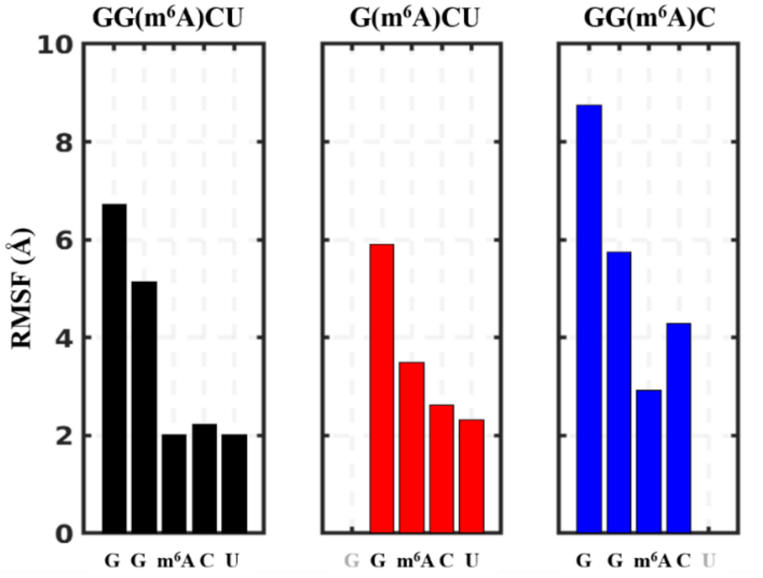

Supplementary Figure S4. Analysis of key interactions between YTHDC1 and m$^6$A. (A) Hydrogen bond interaction distances of the m$^6$A base with the YTHDC1 binding pocket. The interaction distance was measured between two heteroatoms corresponding to acceptor and donor atoms for defining a hydrogen bond. Three interaction distances were measured, namely 1: Ser378(O)--m$^6$A(N6), 2: Asn367(ND2)--m$^6$A(N1) and 3: Asn363(N)--m$^6$A(N3). The bar indicates the average distance value calculated on all configurations collected from 5 independent trajectories for a specific system. The red error bar denotes the standard deviation of the distance. The detailed interaction modes for these hydrogen bonds are shown in the 3D structure. (B) Time series of hydrogen bond interaction distances. The time series of five replicas for a specific system are separated by dashed lines.

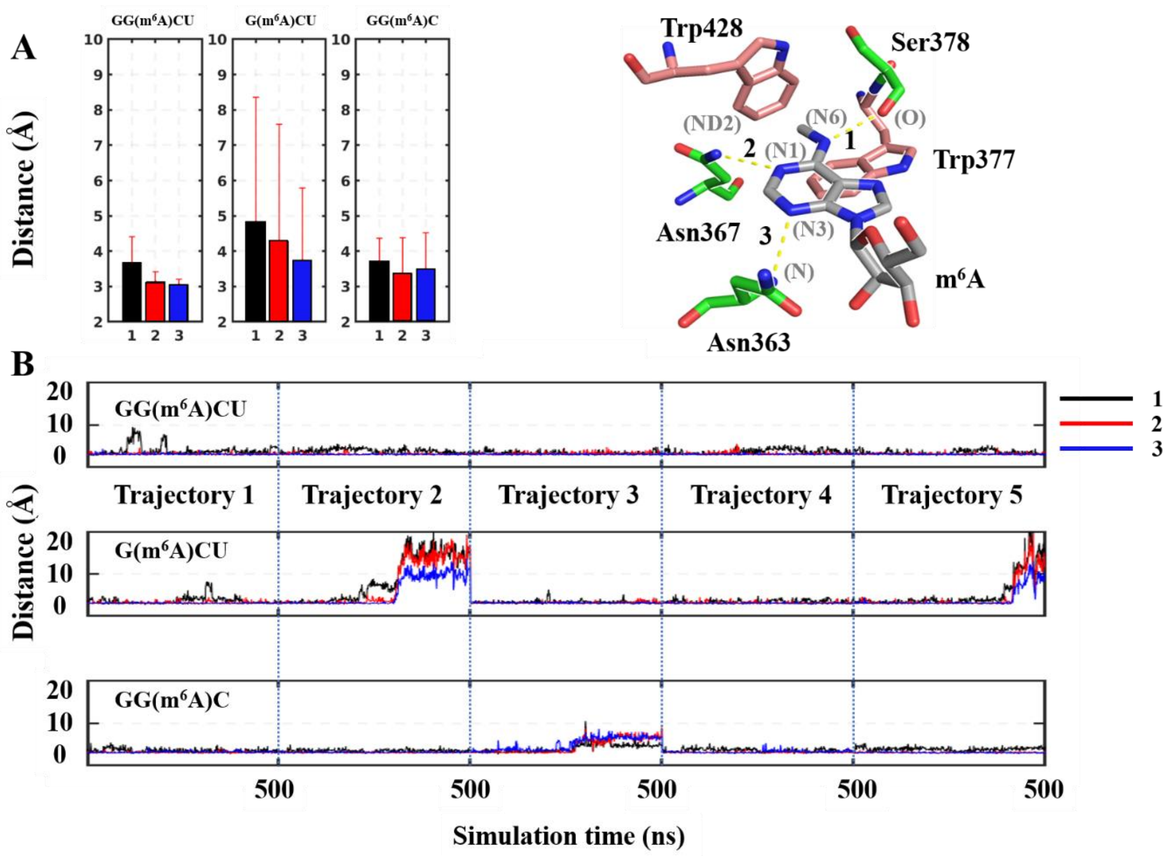

Supplementary Figure S5. RMSD profile of RNA oligomers in the bound state. The plots for GG(m$^6$A)CU and GG(m$^6$A)C systems identical as those in Figure 4. For the plot of G(m$^6$A)CU (middle panel), the snapshots related to the dissociation of m$^6$A from the YTHDC1 binding pocket are neglected. An 8 Å of RMSD cutoff is used for distinguishing the dissociation snapshots of m$^6$A from its binding snapshots.

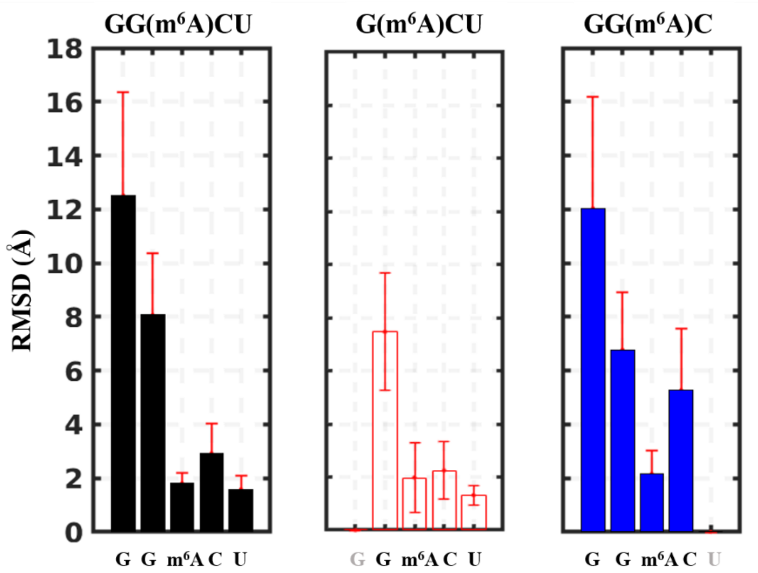

Supplementary Figure S6. Distribution of distances between Arg475 and nucleotide $C_{+1}$ in three different RNA oligomers. The distance was measured between the atom CZ in Arg475 and the geometrical center of the $C_{+1}$ base.

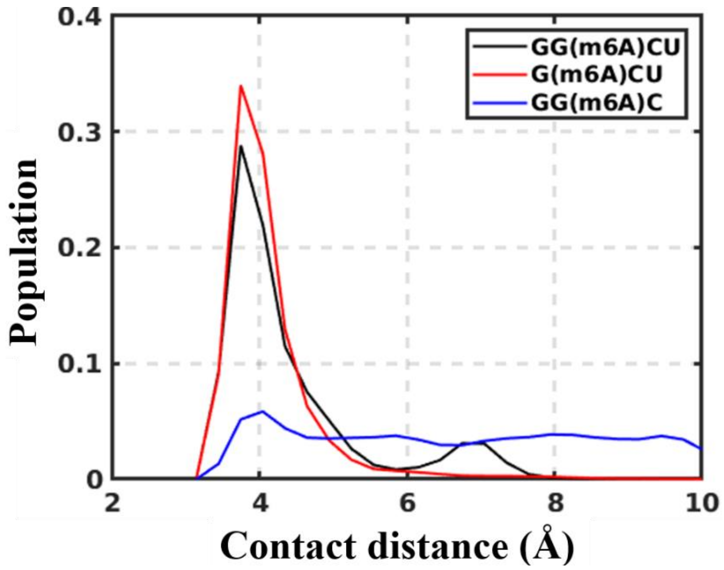

Supplementary Figure S7. Distribution of contact distances for pairwise nucleotides of m6A-containing RNA oligomers in the unbound state. Four pair of contact distances are described by probability distribution plots (indicated by dashed double sided arrows in the structure). The contact distance is calculated in the same manner as that for contact maps.

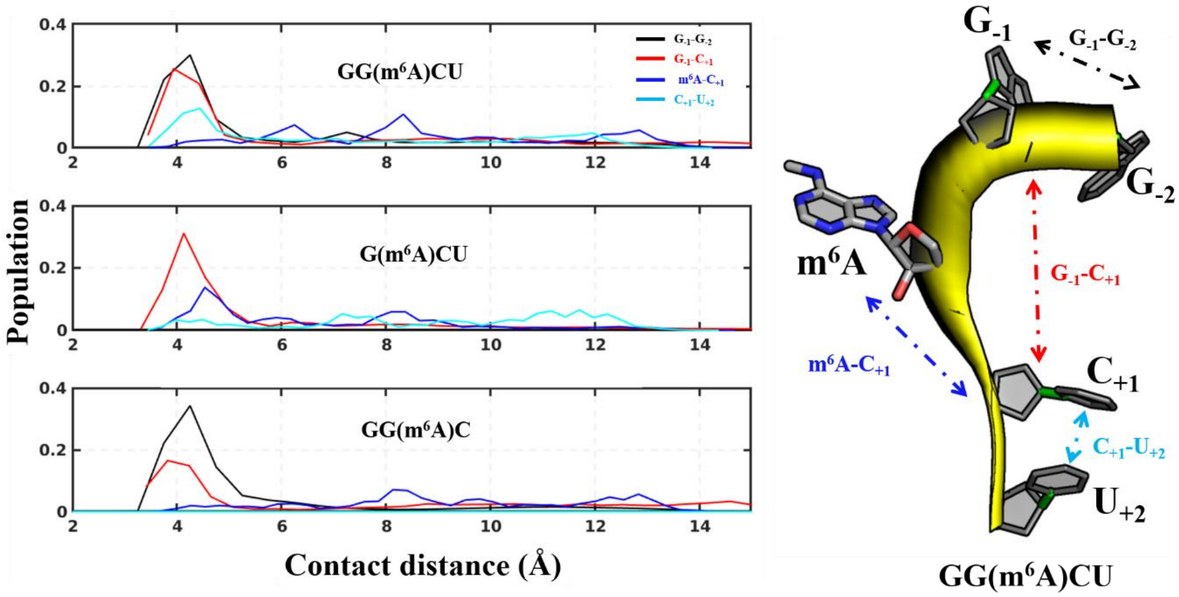

Supplementary Figure S8. Distribution of solvent accessible surface areas (SASA) of (A) methylated adenosine (m$^6$A) and (B) unmethylated adenosine, and (C) conformational landscapes of methylated and unmethylated GGACU in their unbound states. Only the bases of m$^6$A and adenosine on different RNA oligomers are involved in the SASA calculation. Four ligands, viz., GG(m$^6$A)CU, G(m$^6$A)CU, GG(m$^6$A)C and GGACU (shown in black, red, blue and dashed black, respectively) were simulated in their free states in aqueous solution. The distribution for the simulations of GG(m$^6$A)CU in its complex with YTHDC1 is also shown (green) as a basis of comparison; the atoms of YTHDC1 were neglected for the SASA calculation of m$^6$A in the bound state. The PMF plots were produced in the same way as those in Figure 6.

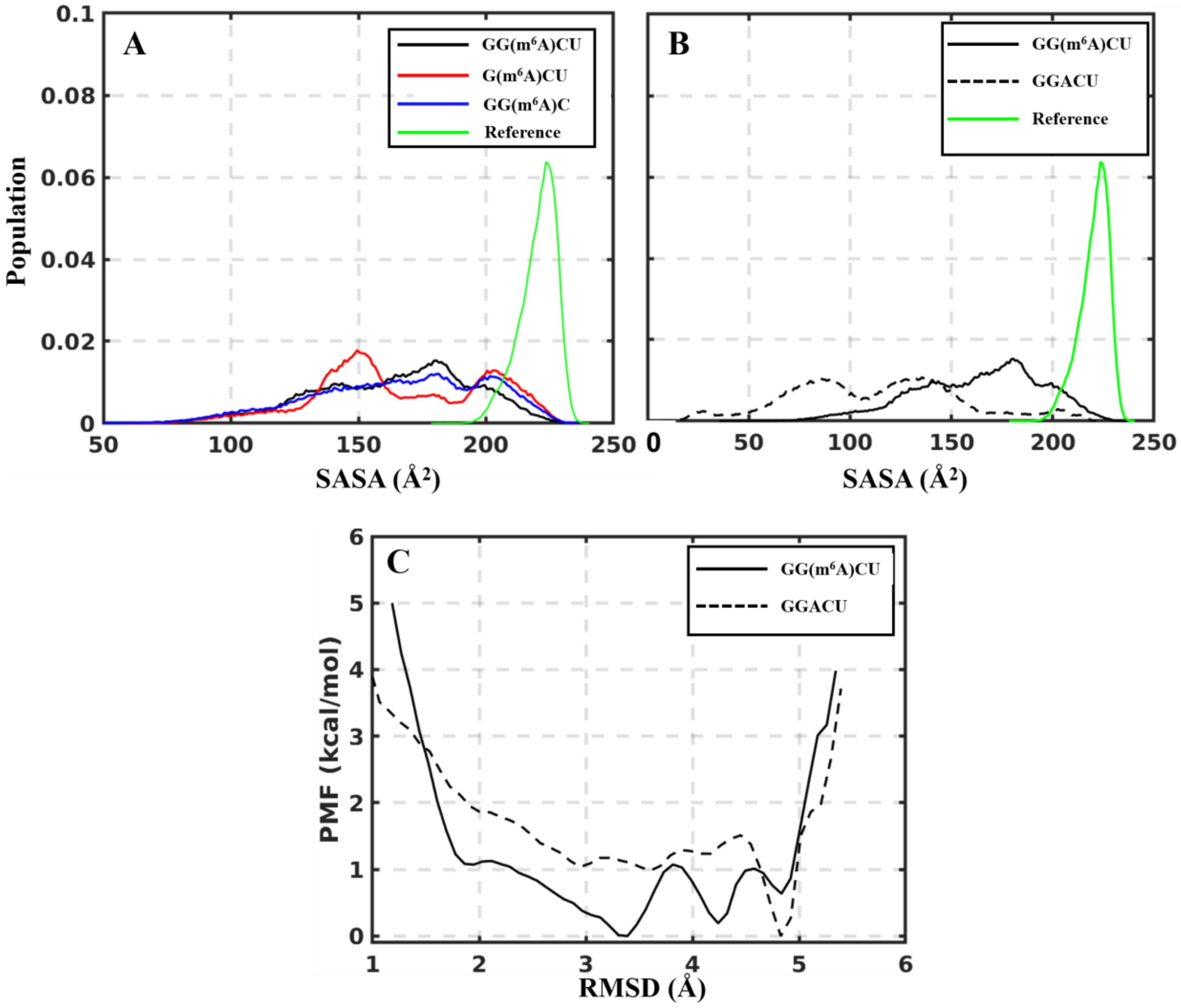

Supplementary Figure S9. Conformational free-energy differences between bound-like and unbound conformations of RNA oligomers in aqueous solution. The RMSD of RNA oligomers with respect to the reference is calculated in the same manner as that used for the PMF plots. The conformational free-energy difference is calculated by the equation $\Delta G = -k_B T ln \frac{p_b}{p_u}$, where $k_B$ is Boltzmann factor, $T$ is temperature (300 K in this study), $p_b$ is the population of bound-like conformations, and $p_u$ is the population of the unbound conformations ($p_u = 1 - p_b$). The value of $\Delta G$ depends on the choice of the RMSD threshold for defining the bound state. The relative $\Delta G$ values for G(m$^6$A)CU and GG(m$^6$A)C are calculated by subtracting the reference values of GG(m$^6$A)CU (dashed lines).

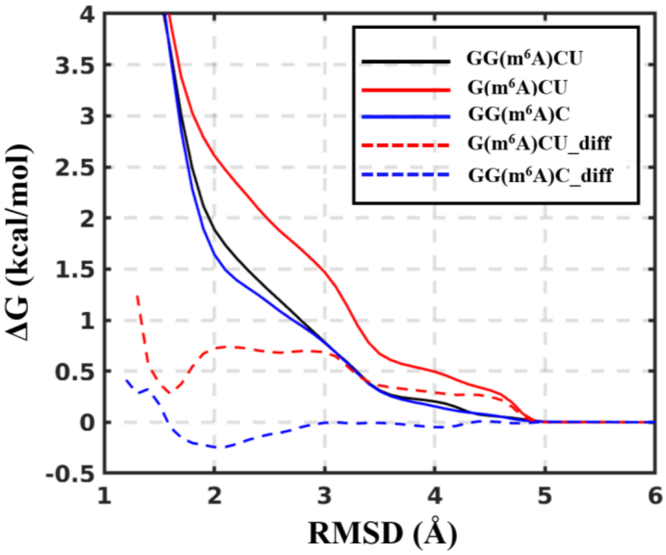

Supplementary Figure S10. Structural comparison between holo YTHDC1 and YTHDF1. (A) YTHDC1 bound to GG(m$^6$A)CU (PDB ID: 4R3I). YTHDC1 shares most key residues with YTHDC2 in the binding pocket. (B) YTHDF1 bound to GG(m$^6$A)CU (PDB ID: 4RCJ). YTHDF1, YTHDF2 and YTHDF3 share all key residues in their binding pockets. Here, the two structures are placed in the same orientation and the corresponding residues are shown in same color schemes.

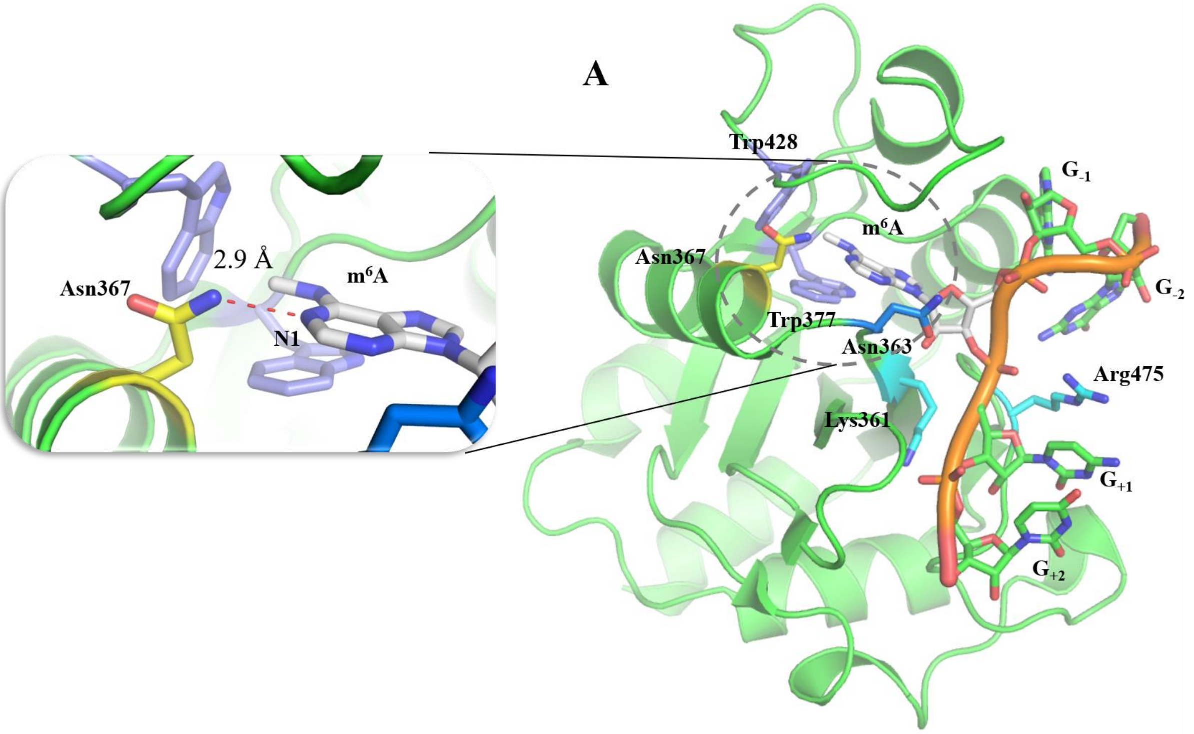

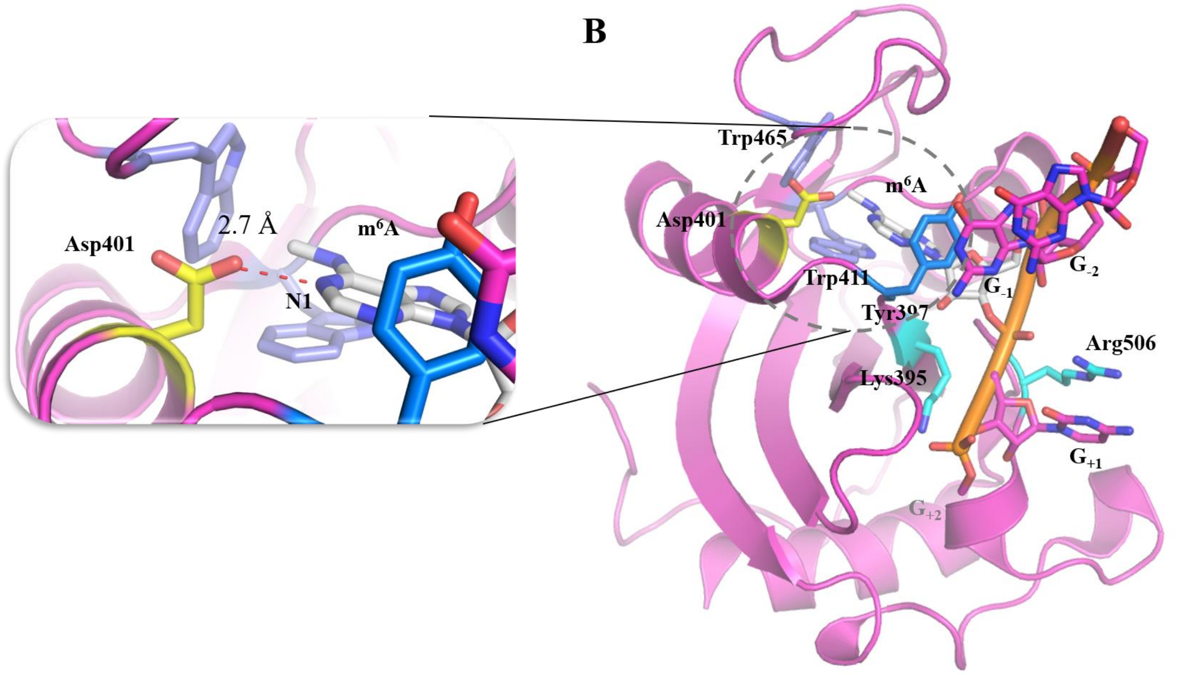

Supplementary Figure S11. Electrostatic potential at the molecular surface of five human m$^6$A-reader proteins. The electrostatic potential (blue, positive; red, negative) was calculated by adaptive Poisson-Boltzmann solver (APBS) electrostatic plugin (version 2.1) within PyMOL. The structures of YTHDC1, YTHDC2, YTHDF1, YTHDF2, and YTHDF3 correspond to 4R3I, 2YU6, 4RCJ, 4RDO, and an in-house structure, respectively. The missing residues were fixed, and hydrogen atoms were generated by CHARMM. The aromatic cage is indicated by the m$^6$A molecule (cyan stick). The two conserved basic residues are highlighted (dashed circles).

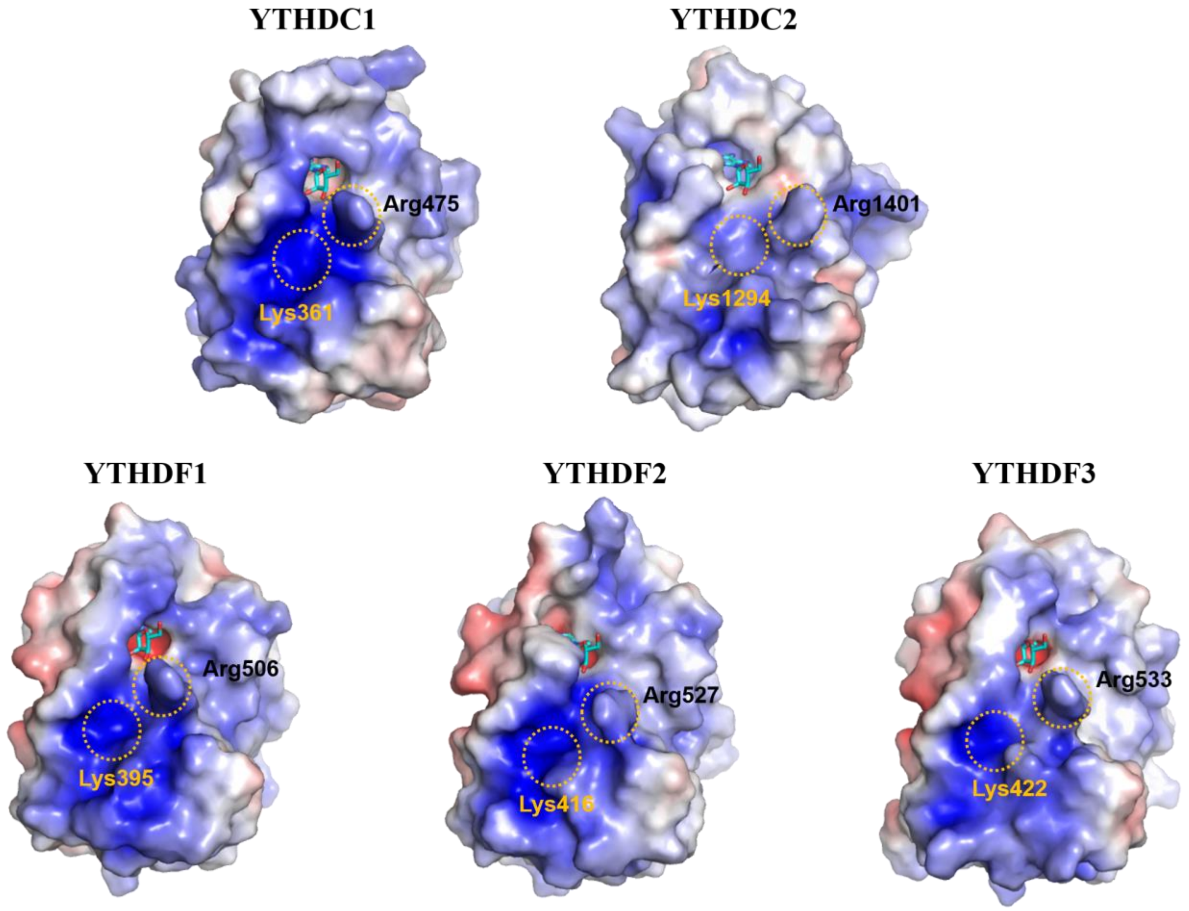
